# A Systematic Benchmark of Machine Learning Methods for Protein-RNA Interaction Prediction

**DOI:** 10.1101/2023.02.14.528560

**Authors:** Marc Horlacher, Giulia Cantini, Julian Hesse, Patrick Schinke, Nicolas Goedert, Shubhankar Londhe, Lambert Moyon, Annalisa Marsico

## Abstract

RNA-binding proteins (RBPs) are central actors of RNA post-transcriptional regulation. Experiments to profile binding sites of RBPs *in vivo* are limited to transcripts expressed in the experimental cell type, creating the need for computational methods to infer missing binding information. While numerous machine-learning based methods have been developed for this task, their use of heterogeneous training and evaluation datasets across different sets of RBPs and CLIP-seq protocols makes a direct comparison of their performance difficult. Here, we compile a set of 37 machine learning (primarily deep learning) methods for *in vivo* RBP-RNA interaction prediction and systematically benchmark a subset of 11 representative methods across hundreds of CLIP-seq datasets and RBPs. Using homogenized sample pre-processing and two negative-class sample generation strategies, we evaluate methods in terms of predictive performance and assess the impact of neural network architectures and input modalities on model performance. We believe that this study will not only enable researchers to choose the optimal prediction method for their tasks at hand, but also aid method developers in developing novel, high-performing methods by introducing a standardized framework for their evaluation.

## 1 Introduction

Out of the over 20,000 annotated human protein-coding genes, at least 1,500 are predicted to code for RNA binding proteins [1]. RNA-binding proteins (RBPs) are involved in a diverse number of functions, such as export and localization of transcripts, post-transcriptional modification, alternative splicing and translation [2] and play an important role in human diseases, such as cancer, neurodegenerative and metabolic diseases [3]. Uncovering the targets of RNA binding proteins is crucial to elucidate their cellular function in health and diseases. Several experimental methods for identifying RBP *in vivo* binding-sites transcriptome-wide have been developed, with arguably the most prevalent being CLIP-seq [4] and its derivatives, such as PAR-CLIP [5], iCLIP [6] and eCLIP [7]. CLIP-seq data is commonly post-processed with peak callers, which identify, from the mapped reads, regions of enriched signal over background, i.e. binding sites. While experimental methods give an unprecedented insight into the binding specificities of RBPs, *in vivo* profiling of protein-RNA interactions is subject to the transcript abundances in the experimental cell type. Thus, researchers must instead rely on computational methods to impute missing binding sites on non-expressed transcripts or to characterize RBP binding sites in settings where no experimental data are available, in order to avoid numerous costly experiments across a wide range of experimental conditions.

Method development for RBP binding-site prediction is an active area of research in the domain of computational RNA biology and an abundance of RBP binding-site prediction methods have been developed in recent years [8, 9, 10]. Development of new methods further accelerated with the advent of deep learning, which showed ground breaking performance improvements in many domains of research, including genomics. Current state-of-the art methods for RBP binding site prediction are usually formulated as a supervised learning problem, to predict whether an RNA sequence is bound or not bound by a certain RBP. Bound regions are usually defined as high-confidence binding sites, so called *peaks*, from CLIP-seq experiments. Models are then trained to classify RNA sequences as bound or unbound, either in a single-task (one RBP per time) or multi-task (several RBP simultaneously) manner [11]. Given this rapid development of several predictive models (1a), it is becoming increasingly difficult for both experimental and computational RNA biologists to select the most appropriate method for the task at hand. This is largely due to the fact that studies train and evaluate their methods on different CLIP-seq datasets, which either encompass a different set of profiled RBPs or may contain binding sites that have been derived via different experimental CLIP-seq protocols (Table 1 and 2). Indeed, it has been shown that different RBPs show a different degree of binding specificity [12] and thus, prediction methods have different upper and lower baselines, depending on the composition of the evaluation dataset. Further, CLIP-seq protocols differ in their signal footprint. For instance, protocols, such as iCLIP [6] or eCLIP [2], profile protein-RNA interaction at single-nucleotide resolution, raising the question whether an increase in predictive performance is due to an improvement in data quality, rather than an improvement of the computational methods. Classification methods require annotation of sequences with *positive* and *negative* labels, such that during training a decision function between the two classes can be learned. While CLIP-seq followed by peak calling explicitly yields a set of positive samples, negative samples are less trivial to obtain. Different negative sample generation strategies were developed across methods, further increasing heterogeneity during method evaluation. To date, multiple studies showed the presence of intrinsic biases in CLIP-seq, such as enhanced crosslinking likelihood at uridines, presence of stick-RBPs and RNase-bias towards termination at guanines [13, 14]. These biases may be predominantly present in the positive class set and may therefore serve as features for class-discrimination, leading to inflated method performances.

**Table 1:** Overview of methods dedicated to predicting the binding propensity of RNA-binding-proteins for a given genomic interval of interest. Star symbol ‘*’ indicates shallow-learning methods. In red are the methods selected for the benchmark.

| Method | Year | Data Type | Evidence | Architecture | Prediction | Ref |
| --- | --- | --- | --- | --- | --- | --- |
| GraphProt* | 2014 | RBP-24 | sequence structure | Graph embedding SVM | Binary | [20] |
| DeepBind | 2015 | RNAcompete | sequence | CNN | Intensity | [45] |
| Deepnet-RBP | 2016 | RBP-24 | sequence structure<br>3D structure | DBN | Binary | [29] |
| iDeepA | 2017 | RBP-24 | sequence | Attention, CNN | Binary | [46] |
| Concise | 2017 | VanNostrand_ENCODE<br>RBP-31 | sequence<br>genomic annotation | spline transformation,<br>CNN | Position-wise<br>signal | [43] |
| iDeep | 2017 | RBP-31 | sequence structure<br>genomic annotation<br>motif, co-binding | DBN, CNN | Binary | [35] |
| iDeepS | 2018 | RBP-31 | sequence structure | CNN, BLSTM | Binary | [30] |
| pysster | 2018 | VanNostrand_ENCODE | sequence | CNN | Binary | [28] |
| DLPRB | 2018 | RNAcompete | sequence structure | CNN,RNN | Intensity | [33] |
| cDeepBind | 2018 | RNAcompete | sequence structure | CNN, LSTM | Intensity | [27] |
| iDeepV | 2018 | RBP-67 | sequence | word2vec, CNN | Binary | [55] |
| iDeepE | 2018 | RBP-24 | sequence | CNN (global+local) | Binary | [47] |
| iDeepM | 2018 | RBP-67 | sequence | CNN, LSTM | Binary | [48] |
| DeepRAM /<br>ECBLSTM | 2019 | RBP-31 | sequence | CNN, BLSTM,<br>k-mer embedding | Binary | [49] |
| SeqWeaver | 2019 | Authors' datasets | sequence | CNN | Binary | [79] |
| RDense | 2019 | RNAcompete | sequence structure | CNN, BLSTM | Intensity | [32] |
| ThermoNet | 2019 | RNAcompete | sequence structure | CNN | Intensity | [74] |
| mmCNN | 2019 | RBP-24 | sequence structure | CNN | Binary | [37] |
| iCapsule | 2019 | Authors' datasets | sequence structure | Capsule Network | Binary | [26] |
| HOCNNLB | 2019 | RBP-31 | sequence | k-mer embedding<br>CNN | Binary | [63] |
| DeepCLIP | 2020 | RBP-24;RNAcompete<br>VanNostrand_ENCODE | sequence | CNN, BLSTM | Binary | [50] |
| DeepRiPe | 2020 | Mukherjee_PARCLIP<br>VanNostrand_ENCODE | sequence<br>genomic annotation | CNN | Multi-label<br>probability | [40] |
| RPI-NET /<br>RNAonGraph | 2020 | RBP-24 | sequence structure | GNN | Binary | [36] |
| DeepRKE | 2020 | RBP-24;RBP-31 | sequence structure | embedding, CNN,<br>BLSTM | Binary | [25] |
| iDeepMV | 2020 | RBP-67 | sequence | Multi-view CNNs<br>ensemble | Binary | [80] |
| DeepA-RBPBS | 2020 | RBP-31 | sequence structure | CNN, biGRU | Binary | [64] |
| MSC-GRU | 2020 | RBP-31 | sequence | CNN, biLSTM | Binary | [65] |
| MultiRBP | 2021 | RNAcompete | sequence structure | CNN | Multi-label<br>intensity | [34] |
| PRISMNet | 2021 | VanNostrand_ENCODE | sequence structure | CNN | Binary | [51] |
| RNAprot | 2021 | RBP-24 | sequence<br>genomic annotation<br>conservation | LSTM | Binary | [42] |
| Multi-resBind | 2021 | Mukherjee_PARCLIP<br>VanNostrand_ENCODE | sequence<br>genomic annotation | CNN | Multi-label<br>probability | [41] |
| RBP-ADDA | 2021 | RNAcompete | sequence | Adversarial domain<br>adaptation | Binary | [81] |
| ResidualBind | 2021 | RNAcompete | sequence | CNN | Intensity | [53] |
| kDeepBind | 2021 | RBP-31 | sequence | k-mer embedding<br>CNN | Binary | [62] |
| RBPSpot | 2021 | ENCORI | sequence<br>structure | embedding<br>DNN | Binary | [54] |
| BERT-RBP | 2022 | VanNostrand_ENCODE | sequence | Language model | Binary | [59] |
| DeepPN | 2022 | RBP-24 | sequence | CNN, GCN | Binary | [82] |

**Table 2:** Overview of datasets used by methods considered in this review. In red are the datasets selected for the benchmark.

| Dataset | Year | First introduced by | Description and numbers | Ref |
| --- | --- | --- | --- | --- |
| RNAcompete | 2013 | RNAcompete | 207 RBPs, across 231 experiments.<br>Total of 241,000 sequences each 38-41 nt length, split into 2 sets: set A (120,326) and set B (121,031). Each sequence has a score from the log odds ratio of intensities of pull-down vs input. | [44] |
| RBP-24 | 2014 | GraphProt | Sites derived from CLIP-seq experiments.<br>24 experiments (23 from Dorina) for 21 RBPs.<br>16 PAR-CLIP, 4 HITS-CLIP, 4 iCLIP. | [20] |
| AURA2 | 2014 | AURA2 | Database of post-transcriptional regulatory interactions in UTR, including binding sites of RBPs. | [83] |
| RBP-31 | 2016 | iONMF | Mix of PAR-CLIP, iCLIP, and HITS-CLIP.<br>31 CLIP experiments for 19 RBPs.<br>Positive controls: derived from positions with high read-counts.<br>Negative controls: positions sampled from genes without interactions from any of the 19 RBPs. | [84] |
| RBP-67 | 2016 | RNAcommender | 67 distinct RBPs, 72,226 UTRs.<br>Total of 502,178 interactions curated from the AURA2 database. | [85] |
| Mukherjee_PARCLIP | 2019 | DeepRiPe | PAR-CLIP experiments for 59 RBPs profiled in the HEK293 cell line. | [39] |
| VanNostrand_ENCODE | 2020 | Van Nostrand et al | 150 RBPs assessed in two cell-types (HepG2, K562), for a total of 223 experiments. | [7] |

In this study, we describe 37 approaches and systematically benchmark 11 RBP binding-site prediction methods on binding sites of 3 large CLIP-seq repositories, comprising a total of 313 unique CLIP-seq experiments across 203 RBPs. Datasets are derived from (a combination of) common CLIP-seq protocols, including iCLIP [6], eCLIP [2] and PAR-CLIP [15]. We develop an uniform data pre-processing and training set construction strategy for all methods, enabling us to evaluate method performance in an unbiased way and allowing us to contrast methods exclusively based on method-intrinsic properties. Notably, we employ two negative-class generation strategies, where one strategy is agnostic to CLIP-seq biases while the other performs bias-aware sampling. We evaluate common properties of methods and their potential impact on model performance, with respect to architecture design choices and input modalities, such as the use of secondary RNA structure as an auxiliary input. Further, we perform cross-evaluation of models for two different cell types, K562 and HepG2, to address the question of whether models trained on CLIP-seq data from one cell type are suitable for prediction on another. Finally we investigate the potential generalization of methods across CLIP-seq protocols, by performing cross-CLIP-seq evaluation where models trained from one protocol are applied onto data from other protocols.

The remainder of this article is structured as follows: First, we give a brief overview over machine learning methods for protein-RNA interaction prediction with a focus on input modalities and deep learning model architectures. Next, we cover benchmarking datasets and their preprocessing, before introducing methods selected for benchmarking. Third, we introduce our benchmark design, summarized in Figure 1b, including train/test splitting, negative sample generation, and evaluation metrics. Finally, we report and critically discuss benchmarking results.

**Figure 1:**
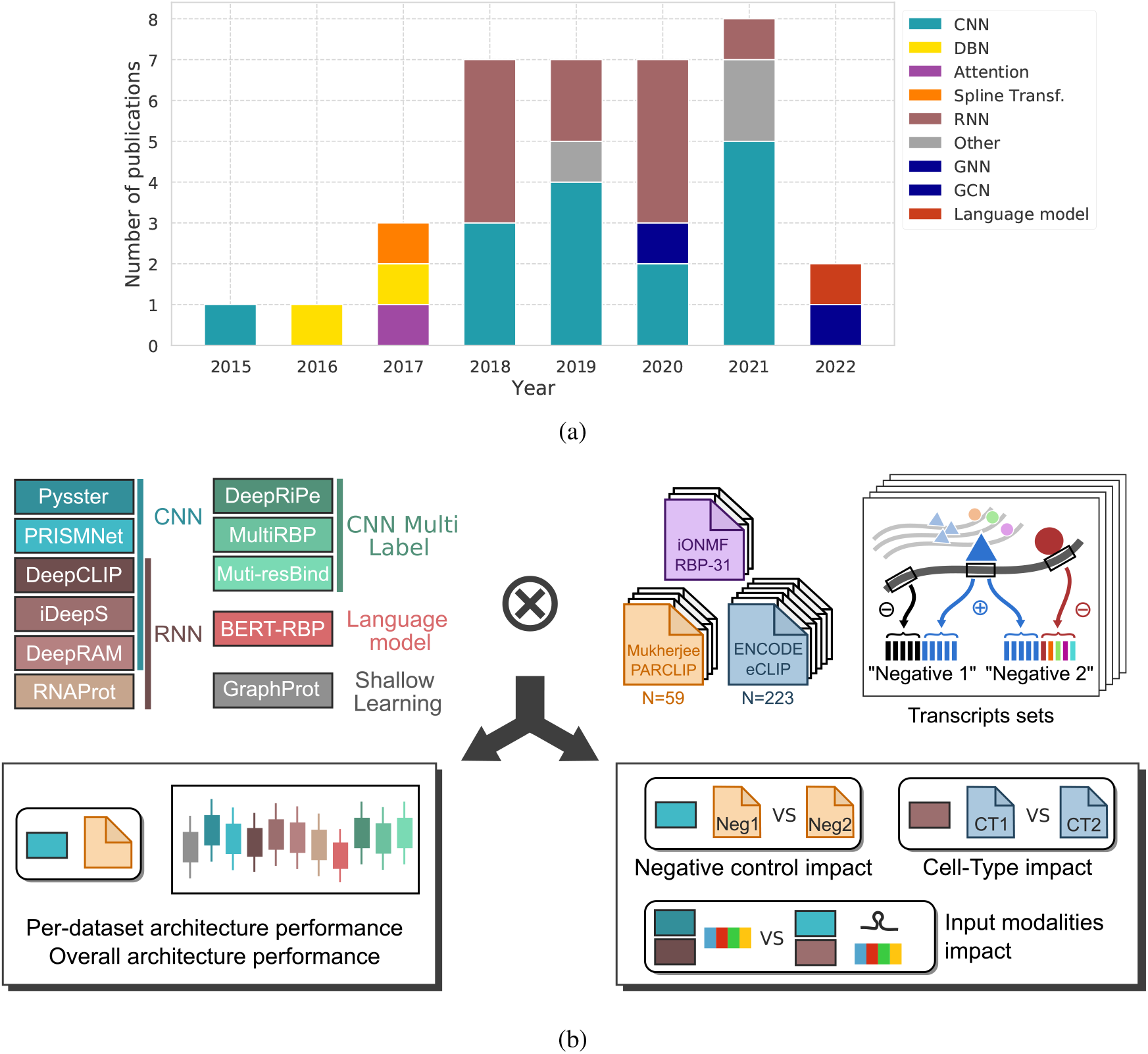
Overview of models and schematic of our benchmark. A. Cumulative barplot of the published methods (see table 1) representing the evolution of architecture choice over the years. B. Illustration of the benchmark presented in this work. Methods representing various architectures were selected. Three datasets of experimentally derived binding sites were preprocessed into folds, separating sequences basing on their transcript assignment. From the selected models and datasets, we first perform an all-against-all evaluation to rank architectures and models (per dataset and across datasets). We also evaluate the impact of negative control sampling on model performance, as well as input modalities. Finally we explore datasets properties, such as cell-type impact for matched RBPs.

## 2 Predicting Protein-RNA Interaction: From Shallow Learning to Deep Learning

Predicting protein binding sites on arbitrary RNA or DNA sequences is a long-standing and unsolved task of computational biology research. While initial methods were predominantly mechanistic programs, often operating via scanning of sequences for a suitable binding site using a position-weight-matrix (PWM) representation of the protein’s target sites [16, 17, 18, 19], the field soon shifted towards more general machine learning methods, which allow for protein binding prediction without first deriving an intermediate PWM representation. These methods were no longer constrained by the representation of binding preferences as fixed-length PWMs, which allowed for modeling of more complex protein-RNA interaction functions and resulted in greater predictive performance compared to classical approaches. For instance, GraphProt [20], a method based on a Support Vector Machine (SVM), uses string-and graph-kernels to encode the primary and secondary structure of an RNA input. While traditional machine learning methods still relied on manual engineering of input features to classify RBP-bound versus unbound sequences, emergence of deep learning methods enabled quasi end-to-end model training. As a result, the research focus shifted from hand-crafting efficient representations of RNA sequence and auxiliary inputs, towards exploration of efficient deep learning architectures and informative input modalities.

In this study, following a comprehensive literature screening, 37 deep learning methods for the prediction of protein-RNA interaction *in vitro* and *in vivo* were identified. Methods were categorized based on their input modalities as well as neural network architecture elements and are summarized in Table 1.

### 2.1 Input Modalities

The main input modality to predictive models of RNA-RBP binding sites is represented by the RNA primary sequence. Sequences of a fixed or variable length surrounding a potential RBP binding sites are converted to either a numerical or one-hot-encoded representation and fed into the model. Besides RNA sequence as the primary input, models make use of a variety of auxiliary inputs, including RNA secondary and tertiary structure, genomic region context, evolutionary conservation and protein co-binding (Table 1).

#### 2.1.1 RNA Secondary and Higher-Order Structure

The higher-order structure of an RNA sequence has been shown to play a vital role in facilitating interactions with the RNA molecule and proteins [21]. For instance, Roquin-1 binds to transcripts via recognition of specific stem loop structures [22] to regulate the post-transcriptional degradation of its targets. Several methods make use of computational RNA folding tools, such as RNAfold [23] or RNAshapes [24], to predict and incorporate secondary RNA structure as additional method inputs. These methods differ significantly with respect to how predicted structures are incorporated into the model. DeepRKE [25], iCapsule [26], cDeepBind [27], pysster [28], Deepnet-RBP [29] and iDeepS [30] first predict a minimum free-energy (MFE) structure via RNAshapes [24], before projecting each position in the input sequence onto an element from *structural vocabulary*, such as hairpin, multi-loop or stem. Subsequently, the structural vocabulary sequence is one-hot encoded. Notably, Deepnet-RBP additionally uses R3DMA [31] to additionally annotate hairpin and internal loop regions with probable tertiary structural motifs. RDense [32], DLPRB [33] and MultiRBP [34] refine this approach by computing the relative frequency of structural elements at each position via RNAplfold [23]. Here, each position in the input encodes secondary structure as a categorical distribution over structural elements. This represents an extension to one-hot encoding, which only takes into account the single MFE structure. Rather than considering structural elements, iDeep [35] uses RNAplfold [23] to predict the unpaired-probability of each position in the input sequence. RPINet [36] and mmCNN [37] operate directly on the predicted base-pairing probability matrix (BPPM). While mmCNN scans the BPPM via 2D convolutional operations, RPINet encodes base-pairing probabilities between input positions as weighted edges in a graph. Lastly, PrismNet is the only method that uses experimentally derived *in vivo* structure via icSHAPE [38], which yields a position-wise score across input sequencing, indicating whether a given position is paired or unpaired. The score vector is concatenated with the one-hot encoded RNA input before being passed to the model.

#### 2.1.2 Genomic Context

Analysis of transcriptome-wide binding site locations showed that many RBPs preferentially bind to specific genomic regions or landmarks [39]. For instance, splicing-associated RBPs predominantly bind at splice junctions and thus information on whether an input sequence is derived from exons, introns or lies at their junction may serve as additional evidence for protein-RNA interaction prediction. DeepRiPe [40], Multi-resBind [41] and iDeep [35] use a region-vocabulary approach to encode the genomic context of a given input sequence. Here, input positions are first annotated with one of 4 (5 in case of iDeep) genomic region types, including CDS, 5’/3’ UTR, introns and exons, followed by one-hot encoding. Using a similar approach, RNAProt [42] maps positions to exon/intron regions, prior to one-hot encoding. Concise [43] computes the distance of each input position to a set of genomic landmarks, including 5’ splice sites, poly-A sites and transcription start sites. Raw distances are transformed to smoothed representations using spline transformations.

#### 2.1.3 Sequence Conservation, Co-Binding and Motifs

Interaction with RBPs determines the fate of transcripts and thus, disruption of RBP target sites may lead to misregulation of post-transcriptional processes and disease. Therefore, RBP target sites are expected to show a higher degree of evolutionary conservation when compared to non-target sites. Leveraging this fact, RNAProt [42] incorporates phastCons and phyloP conservation scores as auxiliary inputs. To leverage prior knowledge, iDeep [35] compute motif-scores for 102 human RBPs using the CISBP-RNA [44] database. Input sequences are additionally annotated with co-binding information using experimental data from other RBPs.

### 2.2 Deep Learning Architectures for Protein-RNA Interaction Prediction

DeepBind [45] was one of the first methods which employed deep learning for the prediction of protein-binding from nucleotide sequences, demonstrating ground-break on both *in vitro* and *in vivo* protein-RNA interaction datasets. DeepBind makes use of a single 1D Convolutional layer, which consists of a set of short, learnable filters that are applied over the input sequence. Over the course of training, the filter-weights are adjusted to yield high activation scores at sequence locations which represent potential binding targets, loosely resembling PWMs scanning of classical methods. Commonly, the outputs of convolutional filters serve as input to downstream layers, such as additional convolutional or recurrent layers, or are directly fed into a linear classifier, which predicts the final binding affinity of the protein of interest for the given RNA sequence. Convolutional neural networks (CNNs) are at the heart of several protein-RNA predictions methods, including iDeep{A,S,E,M} [46, 30, 47, 48], pysster [28], DeepRAM [49], DeepCLIP [50], DeepRiPe [40], MultiRBP [34], PrismNet [51] and Multi-resBind [41], among others (see Table 1). Pysster [28] increases the number of convolutional layers to 3, resulting in a deeper model which can potentially learn more complex binding functions. An additional increase of the input length to 400 (from 101 in case of DeepBind) further increases the receptive field of the model and allows it to consider a broader sequence context around potential binding sites. DeepRiPe [40] uses a single convolutional layer and jointly predicts binding of several RBPs in a multi-task manner, exploiting the fact that many RBPs shared binding preferences and tend to co-bind. This results in an efficient model representation, as convolutional filters are shared across tasks (RBPs), potentially increasing model performance and training stability. Notably, DeepRiPe scans both sequence and genomic region (Section 2.1.2) inputs with separate CNN modules, before joining their outputs for the final classification. A similar approach is used by iDeepS [30] for the independent processing of sequence and secondary structure inputs. PrismNet [51] and Multi-resBind [41] (another multi-task model) further increase network depth via stacking of convolutional layers, while adding residual connections to combat the vanishing gradient problem. PrismNet [51] additionally makes use of a Squeeze-and-Excitation (SE) module [52] to recalibrate outputs of the first convolutional layer. In contrast to DeepRiPe and Multi-resBind, the multi-task method MultiRBP [34] uses convolutional kernels of varying size in the same layer, in order to accommodate binding footprints of different size, across a large number of RBPs. To increase the receptive field of the CNN model, ResidualBind [53] uses several dilated convolutional layers with exponentially increasing dilation coefficient.

Convolutional filters act as (partial) motif detectors, where more complex binding motifs are constructed from the outputs of previous convolutional layers, as the network depth increases. To aggregate the outputs of convolutional layers across the entire sequence, multiple methods make use of recurrent layers, such as long-short-term-memory (LSTM) or gated-recurrent-units (GRU), which enable efficient learning of long-range dependencies. DeepCLIP [50], cDeepBind [27], iDeepS [30], deepRAM’s ECBLSTM model [49] and DeepRKE [25] use a bidirectional LSTM (BLSTM) on top of the outputs of the preceding convolutional layer. While in case of iDeepS, cDeepBind, deepRAM and DeepRKE, the BLSTM output serves as input to a final linear classifier, DeepCLIP directly returns the sum over the BLSTM output as a binding affinity score. Other methods are based on an exclusively recurrent architecture, such as RNAProt [42], which uses an LSTM that directly operates on the RNA sequence. DLPRB [33] combines convolutional and recurrent layers in an alternative way, by concatenating the outputs of a convolutional and a recurrent module, both operating on the RNA sequence, which are then fed into a final linear classifier.

The majority of methods use one-hot encoding to project the RNA sequence into a machine-readable format. However, several methods, including RBPSpot [54], iDeepV [55] and deepRAM [49], use a word2vec [56] model to first learn an embedding of nucleotide 3-mers in an unsupervised manner. During training, k-mers of the input RNA sequence are projected into the word2vec model’s embedding, which then serves as input to subsequent layers. Recently, Transformer-based models emerged as an alternative architecture to CNN-and RNN-based models in fields of natural language processing (NLP) as well as computational biology research [57, 58]. Pre-trained on large corpora of unlabeled data, these models showed ground-breaking performance when fine-tuned on task specific, labeled data. BERT-RBP [59] uses a DNABERT [60] model, pre-trained on a tokenized version of the human genome and fine-tunes it on *in vivo* protein-RNA interaction data. With over 100 million trainable parameters, BERT-RBP represents the largest-capacity deep learning model for protein-RNA interaction prediction evaluated in this study by a large margin. To jointly incorporate RNA sequence and secondary structure graph representations (Section 2.1.1), RPINet [36] uses a modified graph convolutional network (GCN). In each layer, the current node embedding is updated via a graph convolutional operation on the predicted BPP matrix and a convolutional operation along the sequence axis. To obtain binary predictions, the final input embedding is processed by a LSTM layer.

## 3 Material and Methods

### 3.1 Data and Preprocessing

RBP-binding prediction methods were trained and evaluated on binding sites from three distinct sets of experiments, derived from common CLIP-seq protocols, including eCLIP [2], PAR-CLIP [15] and iCLIP [6]. While these datasets were used as training and evaluation sets for some of the benchmarked methods in this study, no study systematically evaluated their method on all three datasets. The RBP / dataset matrix is shown in Table 2.

#### 3.1.1 ENCODE (eCLIP)

The ENCODE Project [7] contains the largest collection of CLIP-seq datasets to date, encompassing 223 eCLIP [2] experiments for 150 RBPs across two cell lines, HepG2 and K562. It has been utilized by a number of studies for model training and evaluation, including Pysster [28], DeepRiPe [40], DeepCLIP [50], Concise [43], Multi-resBind [41] and PrismNet [51]. For each experiment, a set of high-confidence peaks was obtained by processing the narrow-peaks BED files as follows. First, peaks of both replicates are intersected with transcripts of GENCODE (Version 42) to remove all peaks outside of transcript regions. The remaining peaks of both replicates are then intersected and peaks which are present in only one replicate are discarded. For each peak, we define the base at its 5’ end as the single-nucleotide site of crosslinking between RBP and RNA, as suggested by Dominguez et al. [21]. To reduce the computational burden of the benchmark analysis and to select a set of high-quality cross-linked sites, we select at most the top 20,000 peaks with highest signal fold-change over the size-matched input (SMInput) for each experiment.

#### 3.1.2 iONMF (PAR-CLIP, iCLIP, CLIP, HITS-CLIP)

The iONMF dataset was established by Stražar et al. [61] and has since been used by a number of methods for training and evaluation, including iDeep [35], iDeepS [30], DeepRAM [49], DeepRKE [25], kDeepBind [62], HOCNNLB [63], DeepA-RBPBS [64] and MSC-GRU [65]. It consists of cross-linked sites extracted from 31 CLIP-seq experiments for 19 RBPs and, in contrast to the ENCODE and Mukherjee et al. [39] datasets, it includes data derived from different CLIP-seq protocols, including PAR-CLIP [15], iCLIP [6], and HITS-CLIP [66]. The authors retrieved counts data from the iCount [67] and DoRiNA [68] database and selected, for each experiment, the top 100,000 nucleotide positions with the highest cDNA counts. For positions with a distance of less than 15, only the position with the highest cRNA count was considered while all other positions were ignored, as suggested by König et al. [6]. The authors then sampled at most 10,000 cross-linked sites for each experiment in order to reduce processing time. We obtain train and test sets of the iONMF dataset from github.com/mstrazar/iONMF. After merging of both sets, we discard all negative samples defined as part of the iONMF study to obtain the final set of positive cross-linked sites and perform no further processing.

#### 3.1.3 Mukherjee et al. (PAR-CLIP)

This dataset is a subset of 59 PAR-CLIP experiments in the HEK293 cell line, aggregated from different studies and processed by Mukherjee et al. [39]. It was first used by Ghanbari et al. [40] for training and evaluation of DeepRiPe. Variable-length PARalyzer [69] peak regions in BED format were obtained for each PAR-CLIP experiment and peaks were centered to a single nucleotide “pseudo-crosslink” position, in order to homogenize binding sites with the other datasets. Following Ghanbari et al. [40], we did not lift over genomic positions from GRCh37 to GRCh38, but instead used genome and GENCODE [70] versions for the GRCh37 assembly for all downstream processing of this dataset.

### 3.2 Protein-RNA Interaction Prediction Methods

Among the 37 methods compiled in Table 1, we selected 10 deep learning methods for benchmarking, spanning a wide variety of model architectures and input modalities. As a point of reference for the performance of deep learning methods when compared to traditional machine learning approaches, we included GraphProt [20], a shallow learning method based on Support Vector Machines (SVM).

#### 3.2.1 GraphProt (Steffen et al. 2014)

GraphProt [20] makes use of a SVM together with string-and graph-kernels to incorporate both sequence and predicted secondary structure information for classification of a given RNA sequence as bound/unbound. Specifically, during training, GraphProt selects CLIP-seq peaks of at most 75nt in size, which are extended by 15nt up-and down-stream to yield the model’s *viewpoint*. To improve the quality of secondary structure predictions, the viewpoint is further extended by 150nt up-and down-stream, followed by prediction of the minimum free energy structure via RNAshapes [24]. Prior to feature extraction from the secondary structure graph via an extension of the NSPD kernel [71], additional information on the type of substructures (e.g. stem or hairpin-loop) is added via a hypergraph. Finally, SVM classification is performed on the basis of extracted sequence and graph features. Note that as samples in our benchmark dataset are represented by single-nucleotide positions in the transcriptome as a way to homogenize pre-processing across datasets, we extended each sample up-and down-stream to the maximally allow length of 75+15 nt to create the sample’s viewpoint, which is followed by bi-directional viewpoint extension of 150nt for structure prediction. As GraphProt is an SVM classfier, it does not output positive-class probabilities. To compute auROC performance scores, the classification margin of each test sample (i.e. the distance of the sample to the decision boundary) is used instead.

#### 3.2.2 iDeepS (Pan et al. 2018)

iDeepS [30] takes as input an RNA sequence of 101nt and integrates both sequences and predicted secondary structure information via a bi-modal neural network architecture. The minimum free-energy secondary structure is predicted with RNAshapes [24] to yield a 6-symbol structure alphabet, indicating whether a given position in the input sequence resides in a stem (S), multiloop (M), hairpin (H), internal loop (I), dangling end (T) or dangling start (F). Sequence and structure are one-hot encoded and scanned independently by a single CNN layer. Feature maps of both CNN layers are merged and subsequently scanned by a bi-directional LSTM. Finally, the output feature map is passed through a one-unit linear layer with sigmoid activation, to predict the binding probability of the RBP on a given input sequence.

#### 3.2.3 Pysster (Budach et al. 2018)

Pysster [28] is a Python framework for creating CNN models for genomic sequence-based classification and regression tasks. While Pysster may incorporate additional input features, such as secondary structure or genomic region information, Budach et al. [72] showed that high protein-RNA interaction prediction performance can be achieved using models trained on RNA-sequence alone. Pysster [28] takes as input a 400nt window and scans the one-hot encoded RNA sequence via a stack of 3 convolutional layers. The resulting feature maps then serves as input to a stack of two fully-connected layers. In contrast to other methods, Pysster defines two types of negatives, with one half being sampled uniformly from regions with no overlapping binding sites of the given RBP and the other half being sampled from binding sites of other RBPs in order to combat CLIP-seq specific cross-linking bias. The final output layer then consists of three units (one for the positive class and two for both negative classes) and a softmax activation, assigning predicted probabilities to the three classes. Note that in order to make Pysster predictions comparable to other classification methods, only two (positive and negative) output classes were used in this study and Pysster models were trained on just one type of negative samples. Pysster was trained using early-stopping by monitoring the validation loss on a 85/15 train/validation split for up to 200 epochs with a patience of 15. Following Budach et al. [72], Pysster was trained using 3 CNN layers, each with 150 filters and a kernel size of 18 and otherwise default parameters.

#### 3.2.4 DeepRAM (Trabelsi et al. 2019)

DeepRAM [49] is a tool for building flexible deep learning architectures for binary classification of RNA and DNA sequences. Evaluated against both protein-DNA and protein-RNA interaction prediction datasets, the authors compared several common architectures, including combinations of convolutional, recurrent and embedding layers with respect to their predictive performance on the protein-RNA interaction prediction task. For this benchmark, we selected the best performing architecture, termed *ECBLSTM*, which consists of an RNA sequence 3-mer embedding, followed by a convolutional and bi-directional LSTM layer. As the authors evaluated their ECBLSTM model on the iONMF dataset (Stražar et al. [61]) with a sequence length of 101nt, we generated inputs of similar length across all benchmark datasets. DeepRAM first performs hyperparameter selection by training 40 models and evaluating their performance on 3-fold cross-validation, before selecting the best performing hyperparameters for training of a final model. As the training of 40 models across all benchmark datasets was computationally infeasible, we reduced the number of sampled hyperparameters to 5 when benchmarking of deepRAM’s ECBLSTM architecture. Further, the authors additionally train 5 models using the best hyperparameter configuration on the full set of samples and return the model with the lowest training loss as the final model. This is not expected to significantly impact generalization performance, we omitted this step by training only a single model on the optimal hyperparameter configuration, in order to further improve the runtime of deepRAM. Besides those changes, deepRAM was trained as described by Trabelsi et al. [49].

#### 3.2.5 DeepRiPe (Ghanbari et al. 2020)

DeepRiPe [40] is a multi-label classifier which operates on two input modalities, sequence and genomic annotation, with the latter consisting of a mapping of each position in the input RNA sequence to one of four genomic regions, CDS, intron, 3’ UTR and 5’ UTR. Sequence and genomic region inputs are one-hot encoded, before being processed independently by a convolutional layer. The feature maps of both layers are subsequently concatenated and processed by a CNN or GRU layer, before being passed to a fully connected and output layer for classification. DeepRiPe is trained on non-overlapping windows from the human transcriptome, which are labeled with one or more RBPs. Crucially, this alleviates the need for defining negative samples explicitly during training, as windows which are positives for a set of RBPs serve as negatives for the rest. DeepRiPe bins RBPs based on the number of observed binding sites in the corresponding PAR-CLIP or eCLIP experiment and trains multiple models, one for each bin. Further, input sizes of 50nt and 150nt are used for PAR-CLIP and eCLIP samples, respectively. Here, we instead train a single DeepRiPe model with a RNA sequence input size of 150nt and 250nt for the genomic region input, for each of the three datasets, to make processing consistent across datasets. In contrast to the authors, we also did not remove experiments with less than 1000 binding sites, to enable direct comparison with other methods. The authors trained models on 80% of the training data, while 10% were used for validation and testing, respectively. Here, a 80/20 train/validation split was performed on our generated training samples (Section 3.3.1) and models were trained for at most 40 epochs and early stopping with a patience of 5.

#### 3.2.6 DeepCLIP (Gronning et al. 2020)

DeepCLIP [50] is a sequence-only classifier, which first one-hot encodes an input RNA sequence, before applying a convolutional layer as a motif extractor. The resulting feature map is subsequently fed into a bi-directional LSTM layer to perform binary classification of RNA sequences. Notably, in addition to classification, DeepCLIP yields a single-nucleotide binding profile along the input sequence, separating it from the majority of tools for protein-RNA interaction prediction, which usually exclusively perform binary classification of a given input sequence. Similar to GraphProt [20] and RNAProt [42], DeepCLIP takes variable-length input sequence and was evaluated on binding site regions of 12–75nt. In this benchmark, all inputs were length-normalized to a 75nt window centered around the RBP binding site. Further, the authors trained DeepCLIP for a varying number of epochs together with early stopping, depending on the size of the given training dataset. As no specific guidelines for determining the maximum number of training epochs as well as the early stopping patience are provided by the authors, the most prominent choice of *max*_*epochs* = 200 and *patience* = 20 from the publication [50] is used for benchmarking DeepCLIP in this study.

#### 3.2.7 PrismNet (Sun et al. 2021)

PrismNet [51] is the first method to incorporate RNA sequence and experimental structure data measured with *in vivo* click selective 2’-hydroxylacylation and profiling experiment (icSHAPE) [38] to predict RBP binding. icSHAPE assigns reactivity scores at transcriptome-wide level and nucleotide resolution. These scores range between 0 and 1, with the lower scores indicating less reactivity and therefore likely representing paired positions. Sun et al. generated icSHAPE data for seven different cell lines, including HepG2, K562 and HEK293, covering both the ENCODE and Mukherjee datasets. The input to the model is a one-hot encoding of a 101nt input RNA sequence, to which the secondary structure icSHAPE-score vector is appended. The concatenated input is fed into a neural network consisting of a squeeze-and-excitation (SE) module and multiple convolutional layers with residual connections. A fully-connected layer then performs the final binary classification of the input to bound/unbound. We obtained icSHAPE data from Sun et al. [51], lifting over coordinates from GRCh38 to GRCh37 for the Mukherjee and iONMF datasets, using UCSC’s LiftOver tool. As no matching icSHAPE was available for the U266 cell line in the iONMF dataset, icSHAPE vectors were defaulted to −1.0, to indicate missing values. As only two experiments were affected (IDs 19 and 20), benchmark evaluation results are not expected to be affected significantly. Training was performed for 200 epochs using earlystopping with a patience of 20 and a training/validation split of 80/20.

#### 3.2.8 MultiRBP (Karin et al. 2021)

MultiRBP [34] is trained on RNAcompete [73] and subsequently evaluated on eCLIP data. It can be trained on sequence as well as structure, but was demonstrated to performed best when being trained only on the former with an input sequence length of 75 nucleotides. Similar to DeepRiPe [40] and Multi-resBind [41], MultiRBP is a multi-task model which predicts binding affinities for multiple RBPs at once. The CNN-based architecture involves two different branches with varying filter size, which are concatenated just before the output layer, in analogy to ThermoNet [74]. Both include global max-pooling, fully-connected layers and convolutional layers with kernels of varying size. Most notably, since training is done on *in vitro* data, the model is trained on predicting scalar binding intensities rather than binary labels and evaluated on *in vivo* classification without adaptation of the model output. Here, we trained MultiRBP without changing the implementation of the model apart from the size of the output vector, in order to match the amount of RBPs in the three datasets. Data was preprocessed in analogy to DeepRiPe, but with an input size of 75nt. As described by the authors, training was performed for 78 epochs without early stopping on a validation set.

#### 3.2.9 RNAProt (Uhl et al. 2021)

RNAProt [42] is a toolkit for RBP binding sites prediction, integrating a set of utilities from training-dataset generation to reporting statistics and visual information such as logos of extracted motifs. The model is based on RNNs and in its basic configuration takes as input a 81nt RNA sequence, allowing the user to optionally provide additional features, including secondary structure information, conservation scores, exon-intron annotation, transcript region and repeat region annotation. Here we used the default architecture variant (RNN-GRU) and trained the model with a configuration that was performing best according to the benchmark provided by the authors. This configuration makes use of sequence, exon-intron annotation and phyloP [75] and phastCons [76] conservation scores computed on alignments of 100 vertebrates. Following the authors, the model was trained for a maximum of 200 epochs, with early stopping set at 30, and a train-validation split of 80/20.

#### 3.2.10 BERT-RBP (Yamada et al. 2021)

BERT-RBP [59] is a sequence-based model that makes use of DNABERT [60], a large nucleotide language model pre-trained on the human genome, which is fine-tuned to perform protein-RNA interaction binary classification. Given a 101nt input RNA sequence, the sequence is first tokenized into overlapping 3-mer nucleotides. Tokens are embedded and then fed into a 12 layer transformer encoder. The final encoding of the *CLS* token, prepended to the RNA sequence input, is then passed to a classifier, in order to predict binding/non-binding. Following instructions in the author’s Supplementary Table S1, BERT-RBP was trained for 5 epochs. With over 100 million trainable parameters, BERT-RBP represents the largest model evaluated in this benchmark.

#### 3.2.11 Multi-resBind (Zhao et al. 2021)

Multi-resBind [41], similarly to DeepRiPe, trains a multi-task model on CLIP data, but uses a deeper architecture, adding more convolutional layers and residual skip connections. The model was trained on all possible combinations of sequence, structure and region features, but performed best with only 150nt long sequence and region features as a concatenated input vector. We used the exact same preprocessing routine as in DeepRiPe with the exception that we extracted region features with a size of 150 rather than 250. This was done since Multi-resBind, as opposed to DeepRiPe, is a single input model and requires all input features to have the same size. Further steps were done like in the publication, training the model for 40 epochs and evaluating the one with the best validation loss. As done for DeepRiPe, we trained one model on the entire dataset rather 3 different ones as done in the paper in order to keep the training consistent across methods.

### 3.3 Benchmark Design

#### 3.3.1 Transcript-Level Training/Test Splitting

Machine-and Deep-Learning methods require training-and test-sets for the parameter learning and subsequent estimation of the model generalization performance. Crucially, those sets must not intersect in order to avoid over-estimation of the model performance. To this end, several methods randomly split binding sites into training-and test-sets, which may violate the empty-intersection requirement, as peak callers such as CLIPper [77] or PARalyzer [69] can produce overlapping peak regions. Consequently, randomly splitting binding sites into training-and test-sets may lead to over-estimation of model performance.

To ensure that training-and test-sets do not intersect, we employ a transcript-level approach by first dividing human coding and non-coding transcripts into equally-sized, non-overlapping sets and subsequently assigning binding sites to each set. Transcript regions were gathered from GENCODE Version 42, for both the GRCh38 and GRCh37 assemblies, and transcripts overlapping on the same strand were merged and subsequently split into 5 equally-sized sets. Binding sites of each experiment are then intersected with merged transcripts, thus assigning them to one of the 5 sets, such that each set contains roughly 20% of an experiment’s binding sites. While binding sites directly serve as positive-class instances, negative-class instances for a given set were generated exclusively using (merged) transcripts within the set (as described in Section 3.3.2), in order to prevent data-leakage between sets. Evaluation of generalization performance was then performed on the first set, while training was performed on the union of samples of the remaining sets. Notably, this approach allows for a 5-fold cross-validation evaluation of methods, which was omitted due to the computational burden associated with training and evaluation of a large number of deep learning models.

#### 3.3.2 Generation of Negative-Class Samples

Machine learning methods for binary classification require both positive and negative sample instances. However, through CLIP-seq experiments, only positive (i.e. cross-linking) events are observed explicitly, while negatives (i.e. no cross-linking) events need to be defined implicitly, for instance via the absence of observed binding. Several methods, including DeepBind [29], iDeepS [30] and PrismNet [51], sample negatives uniformly from transcriptome regions not overlapping with observed binding sites. This assumes that under absence of an RBP-specific binding features, identifying cross-linked positions is equally (un-)likely across all transcriptome positions. Some studies, including RNAProt [42] and DeepCLIP [50], refine this approach by restricting sampling of negatives to transcripts harboring at least one observed binding site. This ensures that the negatives are derived from transcripts expressed in the experimental cell type and therefore present as a binding partner for the RBP of interest at the time of the experiment, constraining the previous assumption. Recent studies suggest that CLIP-seq experiments suffer from several technical biases, such as background signal, highly abundant RNA, enhanced photoreactivity of uridines or library contamination with RNA fragments of other RBPs [14, 13, 78]. Models trained with negative instances sampled uniformly from unbound regions of (expressed) transcripts are prone to incorporate these biases in their learned function, as CLIP-seq biases are expected to be predominantly present in positive instances. Thus, these models may only partially predict true protein-RNA interaction and performance estimates may therefore be overly optimistic.

To combat this, different strategies have been developed in order to prevent models from learning uninformative biases. For instance, Pysster [28] supplements its set of negatives with cross-linked sites of *other* RBPs, while DeepRiPe [40], a multi-task method, eliminates the need of explicit negatives altogether, as the positive-class instance of one RBP may serve as a negative-class instance of another RBP during model training. To compare the performance of different methods in an unbiased and fair manner, we employed the same negative sample-generation strategy for all methods. Throughout this study, methods are trained and evaluated using two negative-class generation strategies. We construct a set of negatives for each experiment by uniformly sampling positions from transcripts overlapping with at least one binding site of the protein of interest, hereafter referred to as *negative-1*. A second set of negatives is constructed by sampling from binding sites of *other* RBPs experimentally assessed in a given dataset. This ensures that CLIP-seq biases are equally present in the positive and negative set, which renders CLIP-seq biases uninformative with respect to positive/negative class separation and thus prevents learning of CLIP-seq biases during model training. This negative set, consisting of positives of other RBPs, is hereafter referred to as *negative-2*. For both strategies, negatives were generated at a ratio of 1:1 with respect to the number of positive-class instances for each experiment and training and evaluation of all methods was performed separately on both sets of negatives.

#### 3.3.3 Generating Method Inputs

Unless otherwise noted (Section 3.2), we construct method inputs as described by the respective authors.

#### 3.3.4 Performance Evaluation Metrics

Method classification performances are reported as the area under the Receiver Operating Curve (auROC) and Precision-Recall Curve (auPR) as well as the F1 score. As we are benchmarking a wide range of classifiers, including single-and multi-task classifiers, a drawback of the auPR and F1 score metrics is that their baseline performance, i.e. the expected performance of a uninformative random classifier, is subject to the class frequency. These metrics set multi-label classifiers (such as DeepRiPe [40], MultiRBP [34] and Multi-resBind [41]) at a disadvantage, as the frequency of a given label *l* is among *n* samples is 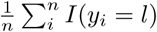 (where *I* is an indicator function) and thus much lower than the positive class frequency of 0.5 in the binary case. Therefore, to evaluate the discriminative power of methods in an unbiased manner, auROC scores were used as the primary evaluation metric in this study.

## 4 Results and Discussion

We trained a total of 4,902 models across a matrix of 313 CLIP-seq experiments and 11 methods in each of the two (negative-1 and negative-2) settings. This includes one shallow learning, 2 and 3 CNN-based and RNN-based binary classification models, respectively, as well as 3 CNN-based multi-task methods (Figure 1).

### 4.1 Deep learning outperforms shallow learning methods

Figure 2a shows the area under Receiver Operating Characteristic curves (auROC) of binary classification methods for models trained on positive and negative-1 samples. Performance of multi-task models (DeepRiPe, Multi-resBind and MultiRBP) is not displayed here, as these methods are trained without universal negative sequences by design. Further, we observed no convergences for 1 deepRAM and 89 BERT-RBP models during training, leading misbehavior during inference (prediction of a single score for all samples) or random baseline performance. These models were removed from the downstream evaluation. As, even after re-training, we observed poor convergence across all BERT-RBP models and could not reproduce the author’s performance, our benchmark results on BERT-RBP are to be treated with care. Consequently, we disregard evaluation results of BERT-RBP in subsequent analysis. GraphProt, the only evaluated shallow learning method, showed the lowest performance with a mean auROC of 0.8211, followed by DeepRAM (0.8697), DeepCLIP (0.8735), RNAProt (0.8857), iDeepS (0.8932), PrismNet (0.9004), while Pysster yielded the highest performance (0.9178) among binary classification methods (Table 3). Figure 2b shows auROC performance in the negative-2 setting. Here, multi-task methods are included, as by design the positives of one RBP may serve as negatives for another. We observe a strong decrease in performance for binary classification method compared to the negative-1 setting (figure 2c), with a mean auROC decrease of −0.0871 (PrismNet), 0.0862 (Pysster), −0.0803(DeepRAM), −0.0786 (DeepCLIP), −0.0761 (iDeepS), and −0.0746 (RNAProt). As outlined in 3.3.2, we hypothesize that this may be due to CLIP-seq experimental biases, which, in case of the negative-1 setting, are exclusively present in the positive sample set, thus serving as a feature for class discrimination. In the negative-2 setting, biases are expected to be equally present in positive and negative samples, making the classification task more challenging.

**Table 3:** Performance of methods as measured by the average auROC from each of the negative-control settings

| **Method** | **Mean auROC**<br>**negative-1** | **Mean auROC**<br>**negative-2** | **Delta auROC** |
| --- | --- | --- | --- |
| BERT-RBP | 0.6681 | 0.5756 | -0.0925 |
| Pysster-101 | 0.8976 | 0.8098 | -0.0878 |
| PRISMNet | 0.9004 | 0.8133 | -0.0871 |
| Pysster | 0.9178 | 0.8315 | -0.0862 |
| DeepRAM | 0.8697 | 0.7894 | -0.0803 |
| DeepCLIP | 0.8735 | 0.7949 | -0.0786 |
| iDeepS | 0.8932 | 0.8171 | -0.0761 |
| RNAProt-Extended | 0.8857 | 0.8112 | -0.0746 |
| GraphProt | 0.8211 | 0.7625 | -0.0586 |
| DeepRiPe | NA | 0.8774 | NA |
| Multi-resBind | NA | 0.8825 | NA |
| MultiRBP | NA | 0.7919 | NA |

**Figure 2:**
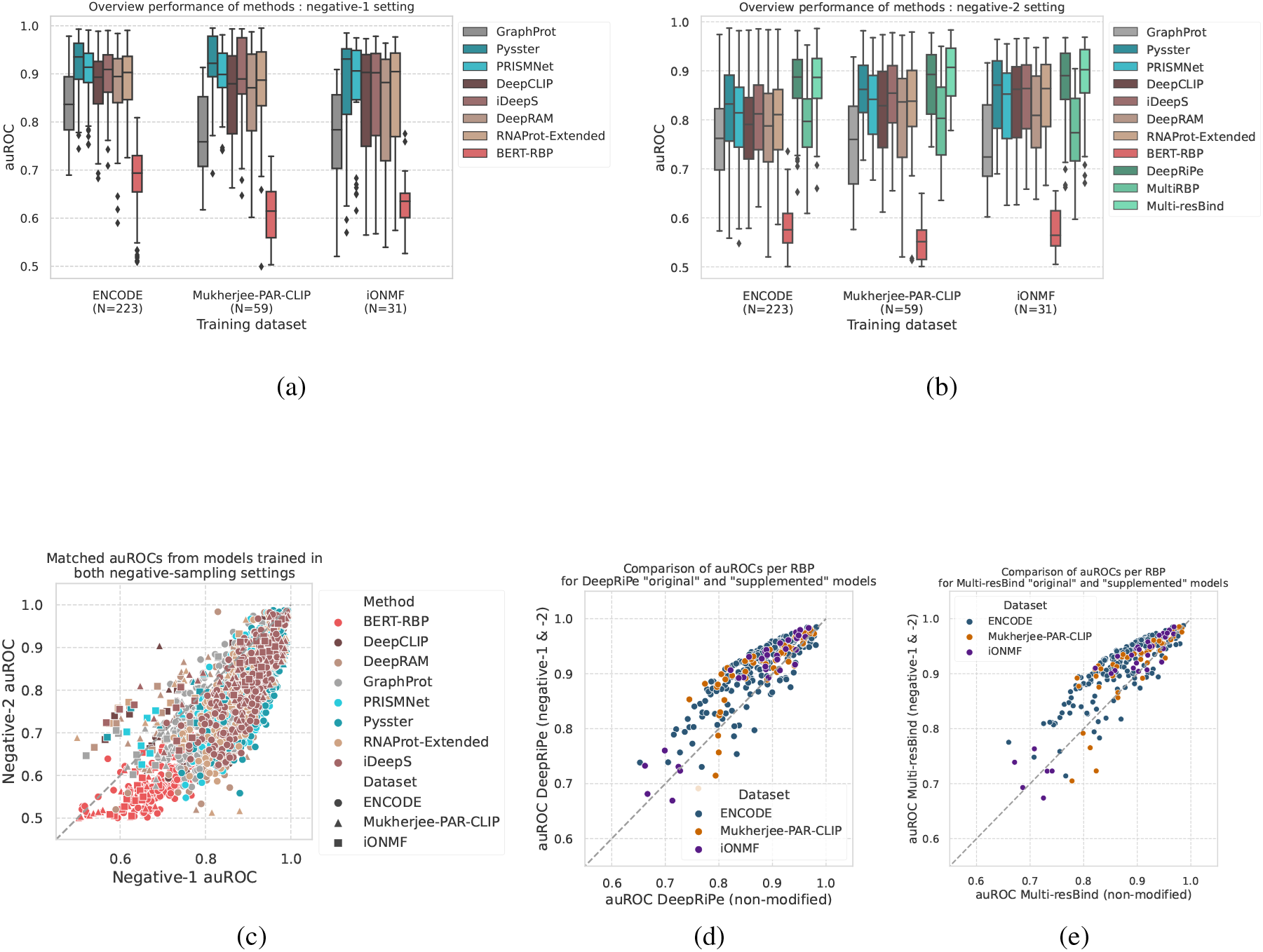
Results from the Benchmark. A. and B. Distribution of auROC values of trained models, across datasets, under the negative-1 (A) and negative-2 (B) sampling schemes. Multi-label models are absent from negative-1 setting due to not handling such negative samples. C. Comparison of auROCs from models trained under the two schemes of negative control sampling. D. and E. Comparison of the classification performance of the modified multi-label methods (enabling negative-1 classification) with the original method, pairing classification results for each RBP, for DeepRiPe in (D) and Multi-resBind (E)

### 4.2 Multi-task outperform binary methods

Except MultiRBP [34], multi-task models outperform binary classification models, with average auROC of 0.8774 and 0.8825 across datasets for DeepRiPe [40] and Multi-resBind [41], respectively. Strikingly, both DeepRiPe and Multi-resBind outperform Pysster, the best binary classification method, by a margin of 0.046 for DeepRiPe and 0.051 for Multi-resBind in auROC performance on the negative-2 setting. The comparably lower performance of MultiRBP (average auROC of 0.7919) may be explained by the fact that this method was initially trained and optimized on *in vitro* RNAcompete data and trained for a fixed number of 78 epochs. Training with the same number of epochs on *in vivo* datasets with a varying number of experiments lead to a considerable degree of over-fitting, as unlike DeepRiPe and Multi-resBind, no early-stopping procedures were used. In addition, MultiRBP operates on a considerably smaller input size of 75nt, compared to 150nt in case of DeepRiPe and Multi-resBind, which may further impact performance. Given the higher performance of binary classification models in the negative-1 setting compared to the negative-2 setting, we speculated that a similar trend may be observed when adding negative-1 to the training of multi-task methods. To this end, we retrained DeepRiPe and Multi-resBind with an additional negative-1 label, by intersecting input tiles with negative-1 sample locations and retaining only those tiles which exclusively overlapped with negative-1 samples (Section 3.2.5 and 3.2.11). Indeed, Figure 2d and 2e show that addition of a universal negative label increases performance of both DeepRiPe and Multi-resBind by and average of 0.0334 and 0.0354, respectively. Note that even with addition of negative-1 labels, multi-task performances are not directly comparable to the binary classification negative-1 setting, as the later makes exclusive use of negative-1 samples for the negative class, which is not the case for multi-task models.

### 4.3 Input size matters

Given that Pysster performed best among binary classification methods, we investigated the impact of Pysster-specific model properties. A distinctive feature of Pysster is its large input size of 400nt, while other methods such a PrismNet, iDeepS and deepRAM operate on a significantly smaller input size of 101nt. To test whether input size is the driver of Pysster’s high performance, we re-trained Pysster across all datasets on an input size of 101nt. Indeed, reducing Pysster’s input size to 101nt lead to a considerable drop of performance (Figure 3a and 3b), such that it now performs on-par with other deep learning binary classification methods such as PrismNet. This effect is maintained when training and evaluating models in the negative-2 setting. Here, we hypothesize that this could due to long-range effects that govern binding of proteins to RNA, such that models benefit from large input windows around potential binding sites. However, low resolution of called peaks across CLIP-seq experiments may also explain this effect, as this could lead to some binding sites not being contained in the model inputs, with an increase in input size alleviating this effect.

**Figure 3:**
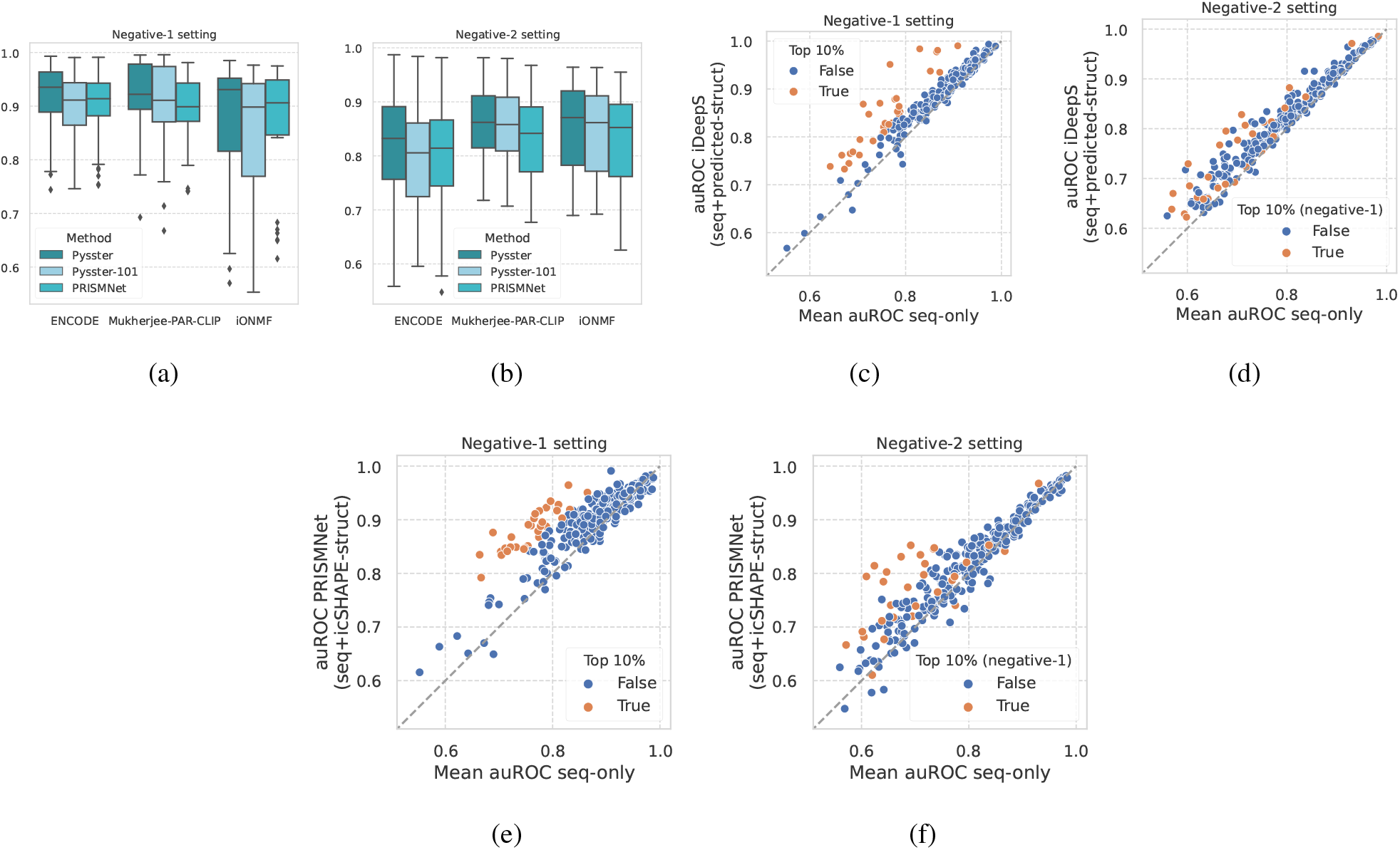
Exploration of methods’ design choice. A. and B. Comparison of Pysster vs “Pysster-101” version where the sequence length of input is reduced from 400 to 101 nucleotides, in the negative-1 setting (A) and negative-2 setting (B). PRISMNet models are included in both for comparison. C. and D. Comparison of the average auROC per RBP from models learning from sequence only (DeepCLIP, DeepRAM, Pysster-101) against auROCs from iDeepS, a model learning from both sequence and predicted structure. Colored are the top 10% RBPs models showing a greater auROC from models learning from both modalities under the negative-1 setting (C). The same models are colored in the negative-2 setting (D) for comparison. E. and F. Same set of plots, here comparing the average auROC per RBP from models learning from sequence only against auROCs from PRISMNet, a method learning from both sequence and *in vivo* measured structure. Colored are the top 10% RBPs models showing a greater auROC from models learning from both modalities under the negative-1 setting (E). The same models are colored in the negative-2 setting (F) for comparison.

### 4.4 Effects of RNA structure as auxiliary input

Given that methods which utilize predicted or experimentally determined RNA structure as auxiliary input did not show a higher performance over sequence-only methods (Figure 2a and 2b), we investigated in more detail whether RNA structure lead to increase performance for individual RBPs. To this end, we compared the performance of structure-aware deep learning methods (iDeepS [30] and PrismNet [51]) with the mean performance of three sequence-only methods (Pysster [28], DeepCLIP [50], and DeepRAM [49]). Note that we used Pysster models trained on 101nt sequence inputs in order to remove any input-size related effects, as iDeepS, PrismNet and deepRAM were trained of sequences of size 101nt. Figures 3c and 3e depict the difference in performance between iDeepS and PrismNet and sequence-only binary classification methods, respectively. While structure does not improve performance for a majority of RBPs, the figures show several outliers for which performance of sequence and structure based methods is elevated above sequence-only methods, including. Examples of RBPs that appear to benefit from structure information include EWSR1, ELAVL2 and CAPRIN1 for which higher performance is observed both for iDeepS and PrismNet. Interestingly, this effect appears to be slightly reduced in the negative-2 setting (Figures 3d and 3f). This may suggest that structural features that guide the discrimination between bound and universally unbound regions are less informative for discrimination of binding sites of two or more RBPs.

### 4.5 Method performance is correlated across CLIP-seq experiments

We next investigated how training and evaluation data affects predictive performance across methods. Figure 4a depicts the auROC performance of each method across all CLIP-seq experiment as a function of the median performance of methods for the given experiment. One can observe that the performance per method across experiments correlates strongly with their median auROC across methods. Notably, the variance in auROC across methods is greater for experiments with overall lower performance as compared to high-performing experiments (bottom half versus top half - Levene statistics = 394.77; p-value < 1.139e-82). This effect is pronounced for the negative-2 setting (Figure 4b), which shows a strong linear correlation between a method’s performance and the median performance across methods for an experiment. This is likely due to different training set sizes across experiments, which vary greatly and have a significant effect on model performance (Figure 4c). As expected, we measured a significant positive correlation of training set size and model performance for both the ENCODE and the PAR-CLIP datasets (ENCODE: Spearman *r* = 0.396, *p <* 1.829*×* 10^−84^; PAR-CLIP: *r* = 0.196, *p <* 1.345*×* 10^−5^), while the iONMF dataset was excluded, as it has the same training set sizes across all experiments.

**Figure 4:**
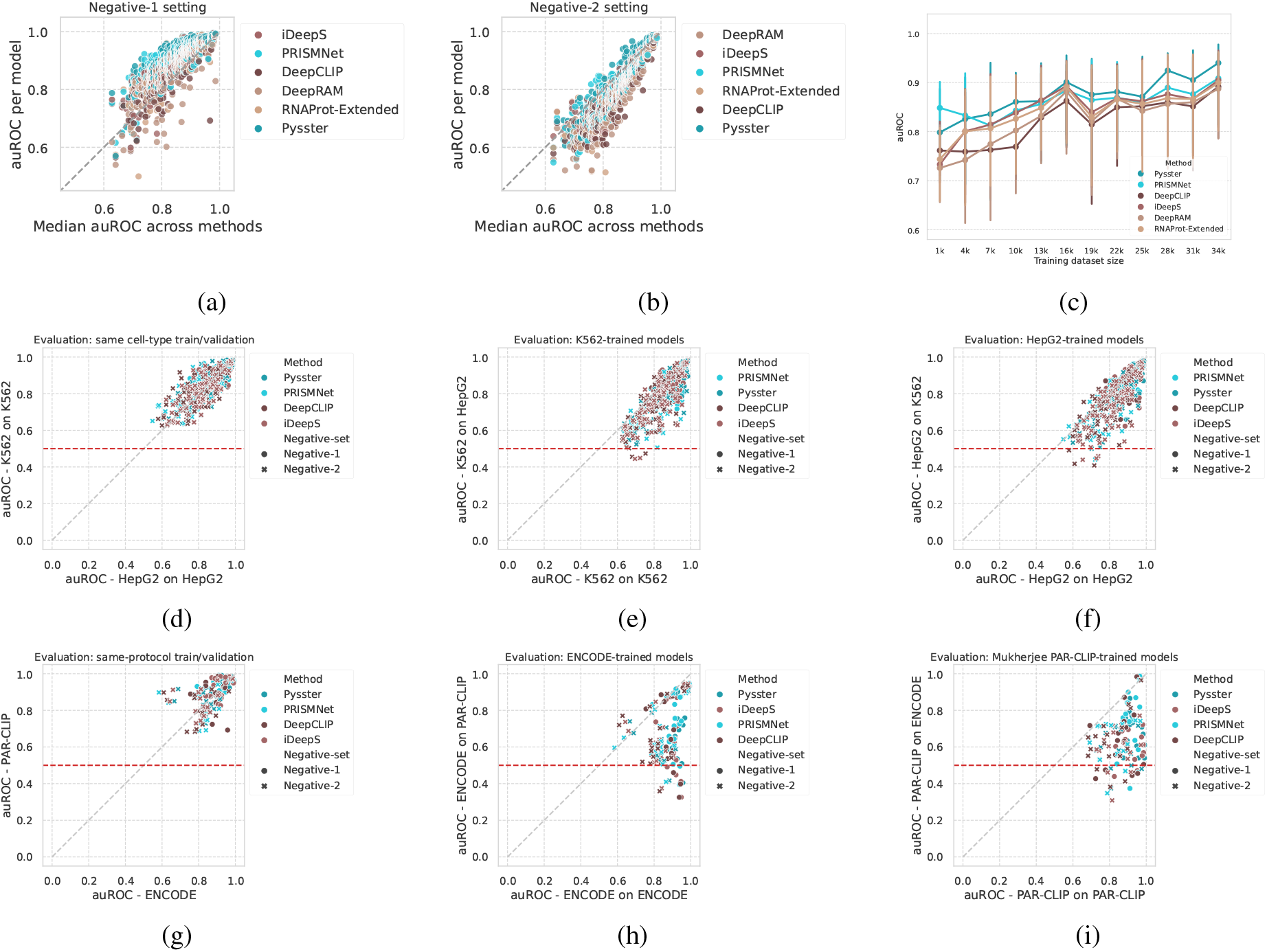
Influence of input modalities. A. and B. Stability of models’ performance per RBP across single-label architectures. Each point is a model for an RBP and a method, plotted against the RBP’s median auROC across methods, in the negative-1 (A) and negative-2 (B) settings. C. Correlation between model performance and dataset size, over the range of dataset sizes from ENCODE. Models are grouped per training dataset bin size (bin size = 2,000). Dots represent the median AUROC of models per bin, for each method. Error bar: 25-75% interquartile. D. Comparison of auROCs for 73 RBPs evaluated in two different cell-types from the ENCODE dataset. Models are paired on the RBP names, while the auROCs are computed on sequences derived from the same-cell type used for training. E. and F. Comparison of auROCs for 73 RBPs evaluated in two different cell-types from the ENCODE dataset, comparing the performance on same-cell-type evaluation (x-axis) against the performance from cross-cell-type evaluation (y-axis) for K562-trained models (E) and HepG2-trained models (F). Red line: random performance. G. Comparison of auROCs for 17 RBPs matched between Mukherjee’s PAR-CLIP and ENCODE eCLIP experiments. Models are paired on the RBP names, while the auROCs are computed on sequences derived from the same experimental-protocol used for training. H. and I. Comparison of auROCs for 17 RBPs matched between Mukherjee’s PAR-CLIP and ENCODE eCLIP experiments, comparing the performance on same-protocol evaluation (x-axis) against the performance from cross-protocol evaluation (y-axis) for ENCODE trained models (H) and PAR-CLIP trained models (I). Red line: random performance.

### 4.6 Models partially learn cell-type specific binding

The ENCODE dataset consists of 223 eCLIP experiments across two cell lines, HepG2 (103) and K562 (120), with 73 RBPs being covered by both cell lines. Figure 4d compares the performance of DeepCLIP, iDeepS, PrismNet and Pysster models across RBPs covered by both ENCODE cell lines. Models did not perform better in one cell line over the other and this effect is consistent over negative-1 and negative-2 samples. We next turned to the question whether models trained on one cell type are applicable (i.e. retain high prediction performance) on another. This is crucial, as a key use case of computational method for protein-RNA interaction prediction is imputation of missing binding information on transcripts no present in the experimental condition, such as unexpressed transcripts. To this end, we selected RBPs covered by both HepG2 and K562 eCLIP experiments and performed cross-predictions, such that models trained on the HepG2 cell line were evaluated on hold-out data from the K562 cell line and vice versa. Figure 4e and 4f show the cross-cell-line performance of model training on the HepG2 and K562 cell lines, respectively. In both cases, a considerable drop in performance is observed, indicating that machine learning models trained on a given cell line learn cell type specific binding feature, which only partially generalize to other cell types. In some cases, models fall to or even below the random baseline auROC performance of 0.5 (red line), although high performing models generally appear to yield high performance even in a cross-cell-line evaluation setting.

### 4.7 Models learn strong protocol-specific biases

We next evaluated whether model performance is subject to the underlying CLIP-seq experimental protocol. To this end, we compared model performances for models trained on RBPs represented by both ENCODE eCLIP and PAR-CLIP experiments from the Mukherjee et al. [39] dataset. While performance differs between protocols for selected RBPs, there appears to be no general trend of better performance on data from one protocol over the other, as shown in Figure 4g. In analogy to our cross-cell-line evaluation, we next evaluated the extent to which models trained on data from one CLIP-seq protocol generalize to data from another protocol. Figure 4h and 4i performance drops significantly when evaluating trained model on data obtained from a different CLIP-seq protocol. We note that besides protocol, ENCODE and Mukherjee et al. make use of different peak callers (CLIPper and PARalizer, respectively), which may impact the final set of binding sites significantly. Further analysis of the impact of peak callers are necessary in order to disentangle the effects of protocol and peak callers on performance drops observed here.

## 5 Conclusion

In this study, we evaluate 11 *in vivo* protein-RNA interaction prediction methods across 313 CLIP-seq datasets with respect to their classification performance on a large cohort of CLIP-seq datasets. Our benchmark revealed that no particular deep learning architecture, such as CNN or RNN, represents a major advantage over others. However, our result revealed that the size of RNA input can have a strong effect on model performance and that multi-task generally outperform single-task methods. We further explored two generation schemes for negative class samples and demonstrated that sampling negatives from unbound regions generally leads to higher performance, possibly due to incorporation of CLIP-seq biases as discriminative features. We demonstrated that predicted and *in vivo* secondary structure might improve model performance for some RBPs, while this effect is subject to the chosen negative samples and is diminished in case negatives are sampled from binding sites of other RBPs. Cross-evaluation results showed that models partially learned cell-type specific RBP-binding, while prediction across protocols leads to a strong decrease in performance, which may be attributed to protocol or peak caller specific biases. We believe that this study will guide future methods development in the field of computational modeling of protein-RNA interaction by serving as a reference for method design in regards to architecture, input modalities and generation of negative controls.

## 6 Data Availability

All data processed in this study was obtained exclusively from public sources (ENCODE [7], Mukherjee et al. [39] and Stražar et al. [61]).

## 7 Code Availability

Code for reproducing the results of this study is available at https://github.com/mhorlacher/Benchmark-RBP.

